# Ennet: construction of potential cancer-driving networks based on somatic enhancer mutations only

**DOI:** 10.1101/216226

**Authors:** Ya Cui, Yiwei Niu, Xueyi Teng, Dan Wang, Huaxia Luo, Peng Zhang, Wei Wu, Shunmin He, Jianjun Luo, Runsheng Chen

**Author notes:** These authors contributed equally to this work (Ya Cui, Yiwei Niu, Xueyi Teng). Correspondence should be addressed R.C.

## Abstract

Whole genome sequencing technology has facilitated the discovery of a large number of somatic mutations in enhancers (SMEs), whereas the utility of SMEs in tumorigenesis has not been fully explored. Here we present Ennet, a method to comprehensively investigate SMEs enriched networks (SME-networks) in cancer by integrating SMEs, enhancer-gene interactions and gene-gene interactions. Using Ennet, we performed a pan-cancer analysis in 2004 samples from 8 cancer types and found many well-known cancer drivers were involved in the SME-networks, including *ESR1*, *SMAD3*, *MYC*, *EGFR*, *BCL2* and *PAX5*. Meanwhile, Ennet also identified many new networks with less characterization but have potentially important roles in cancer, including a large SME-network in medulloblastoma (MB), which contains genes enriched in the glutamate receptor and neural development pathways. Interestingly, SME-networks are specific across cancer types, and the vast majority of the genes identified by Ennet have few mutations in gene bodies. Collectively, our work suggests that using enhancer-only somatic mutations can be an effective way to discover potential cancer-driving networks. Ennet provides a new perspective to explore new mechanisms for tumor progression from SMEs.

## BACKGROUND

Enhancer elements act as regulators of gene expression in biological processes. The Encyclopedia of DNA Elements (ENCODE) Project and other studies have identified millions of enhancers in various tissues [1]. The roles of enhancer in disease have acquired considerable attention. Most germline risk variants identified by GWAS studies reside in putative enhancer elements [2]. Recent studies have also suggested that enhancer malfunction through somatic mutations could be the cause of tumorigenesis by altering the expression of known cancer genes, including *ESR1* [3], *MYC* [4] and *PAX5* [5]. At present, the availability of thousands of whole genomes sequences from tumor samples has paved the avenue for systematic exploration of somatic enhancer mutations.

However, system exploring the role of SMEs in cancer development is far more challenging. Cancer-driving elements can be generally detected by comparing their observed mutation frequency to the expected background rate from this region. Such efforts have leaded to the discovery of a lot of recurrent mutations in the gene body and promoter regions of genes, but failed to identify many recurrently mutated enhancers, even in two large cohorts whole-genome sequencing studies (560 breast cancer and 150 CLL cases, respectively) [5,6]. These results may be due to two reasons: (1) The frequency of mutations occurring in enhancers is too low. Unbiased system analysis of mutations in the whole genome can easily pick up mutational hot spots in gene bodies or promoters, but not in enhancers [7]. (2) The size of whole-genome sequenced samples in specific cancer types are still too small. Approaches that increase samples size by including various types of tumors are effective for gene bodies and promoters, but are not suitable for enhancers, which function in a tissue-specific manner. Hence, new approaches are urgently needed to identify causal enhancer mutations and their downstream consequences.

Knowledge-based network strategies, which assume that complex disease is not caused by individual mutations, nor by single genes, but by combinations of genes acting in molecular networks, have been proved to be an effective way for solving the problems of “sparse mutations” and “small sample size” [8–10]. However, these strategies are difficult to apply to noncoding regulatory elements owing to defective interactions between gene-gene interaction networks and the noncoding elements. Recently, chromosome conformation capture (3C) derived technologies, such as Hi-C [11], Capture Hi-C [12,13], ChIA-PET [14], and many computational methodologies [15–20] offered a lot of ways to appraise and identify the interactions between regulatory elements and genes. These interactions provide us with an opportunity to explore the function of sparse mutations in enhancers with a reasonable sample sizes based on knowledge-based networks.

Here, we present Ennet, a new framework to investigate biological networks that are enriched with somatic enhancer mutations in cancer. Our results derived from 8 cancer types show that exploiting SMEs only can be an effective way to discover potential cancer-driving networks when enhancer-gene interactions are integrated with functional molecular networks. More interestingly, almost none of the genes identified by Ennet, whether known or new, were significantly mutated in their body or promoter regions. It means that these genes may be ignored by the classical gene body or promoter focused methods. Collectively, Ennet provides a new possibility for finding new mechanisms in tumor development.

## RESULTS

### Overview of Ennet

Enhancers are important functional elements that are scattered across chromosomes and regulate the genes through acting on their promoters (Fig. 1A). However, identifying cancer driver genes by using only the mutation information in enhancers is difficult owing to low mutational density in enhancers (Fig. 1C). Thus, we devised Ennet, a new method that integrates the mutation profiles of enhancers, enhancer-promoter interactions and knowledge-based gene networks (Fig 1D). Unlike conventional cancer driver classifiers, Ennet only considers mutations occurring in the enhancers, and the main purpose of Ennet is to identify SME-networks. Ennet begins by counting mutations falling within tissue-specific enhancers of each gene. Then all mutations are summed up across all patients and the counts are mapped to a knowledge-based gene network. After network smoothing (this step is optional), Ennet selects subnetworks with a substantial load of SMEs as potential cancer-driving subnetworks based on the mutational score. Beside identifying SMEs enriched subnetworks, Ennet can also pick out a great quantity of potentially new driver enhancers with very low-frequency mutations that might have important roles in cancer. The functional impacts of mutations on transcription factor binding and target gene expression will require further investigations.

**Figure 1.**
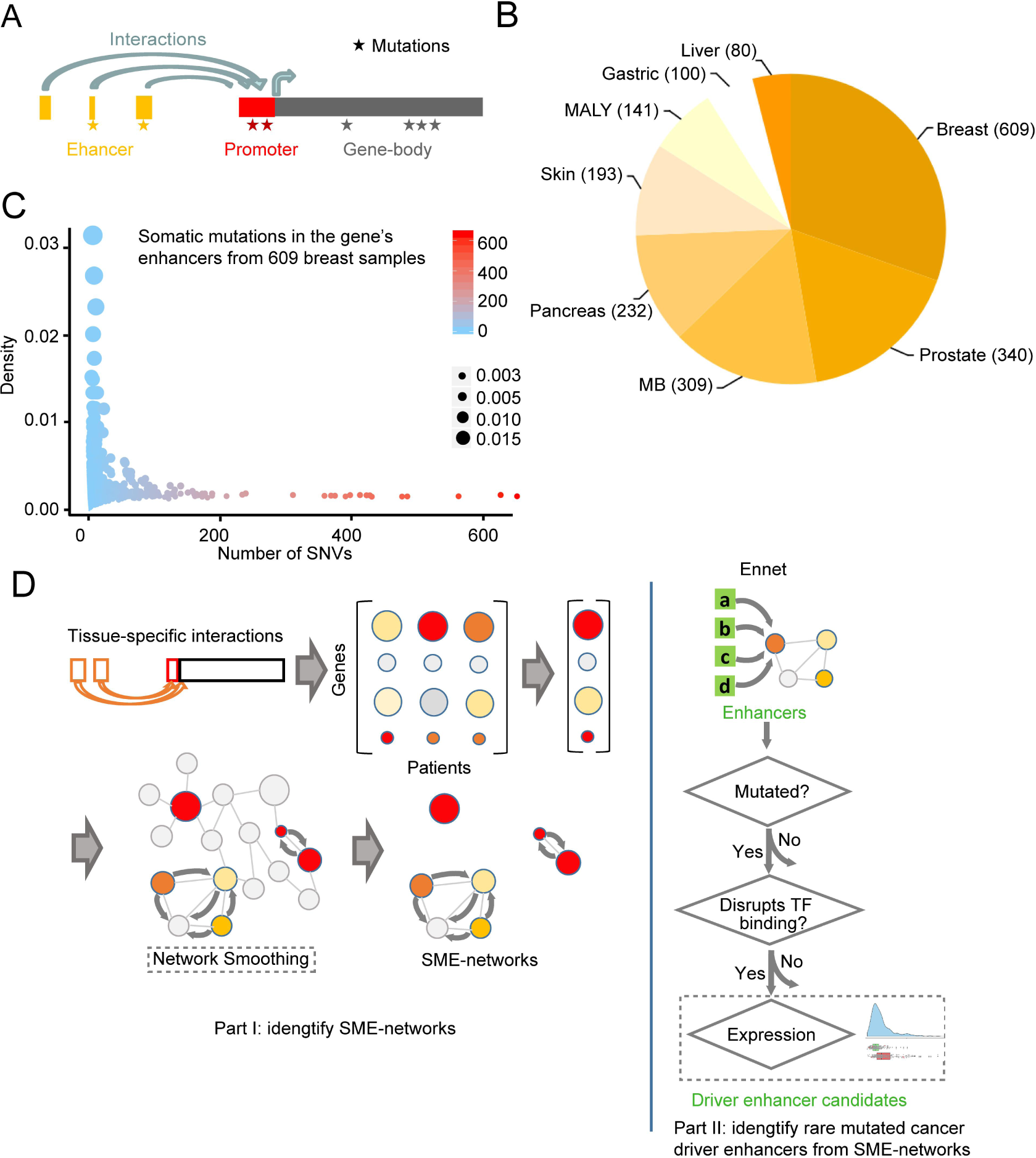
Overview of Ennet. (A) Diagram of gene functional regions: enhancer, promoter and gene body. (B) Sample numbers of the 8 cancer types analysed (MALY: malignant lymphoma, MB: medulloblastoma). (C) Somatic mutation density of genes’ enhancers from 609 breast cancer samples. (D) Overview of the methodology of Ennet. The steps with dashed lines are optional.

### Overview of SME-networks identified by Ennet

We used Ennet to perform an analysis of somatic mutations from 2,004 samples in 8 cancer types (Fig. 1B). Several SME-networks containing altogether 436 genes were identified by Ennet in different cancers (node number > 3 are shown in Fig. 2). Compared these genes with the 125 Mut-driver genes defined by the 20/20 rule [21], 18 genes were found in both collections (Supplemental Figure 1). Moreover, Ennet identified known cancer driver genes such as *EGFR* [22], *SMAD3* [23] and *YWHAZ* [24] as high node degree genes. Importantly, *PAX5* [5] and *ESR1* [3], whose enhancer mutations have been recently reported to promote tumorigenesis, were found by Ennet. A number of well-known cancer pathways such as the *EGFR*, *BCL2*, and *MYC* signaling pathways were also identified by Ennet. Our data indicate that genes identified by Ennet perform the function in cancer-related pathways.

**Figure 2.**
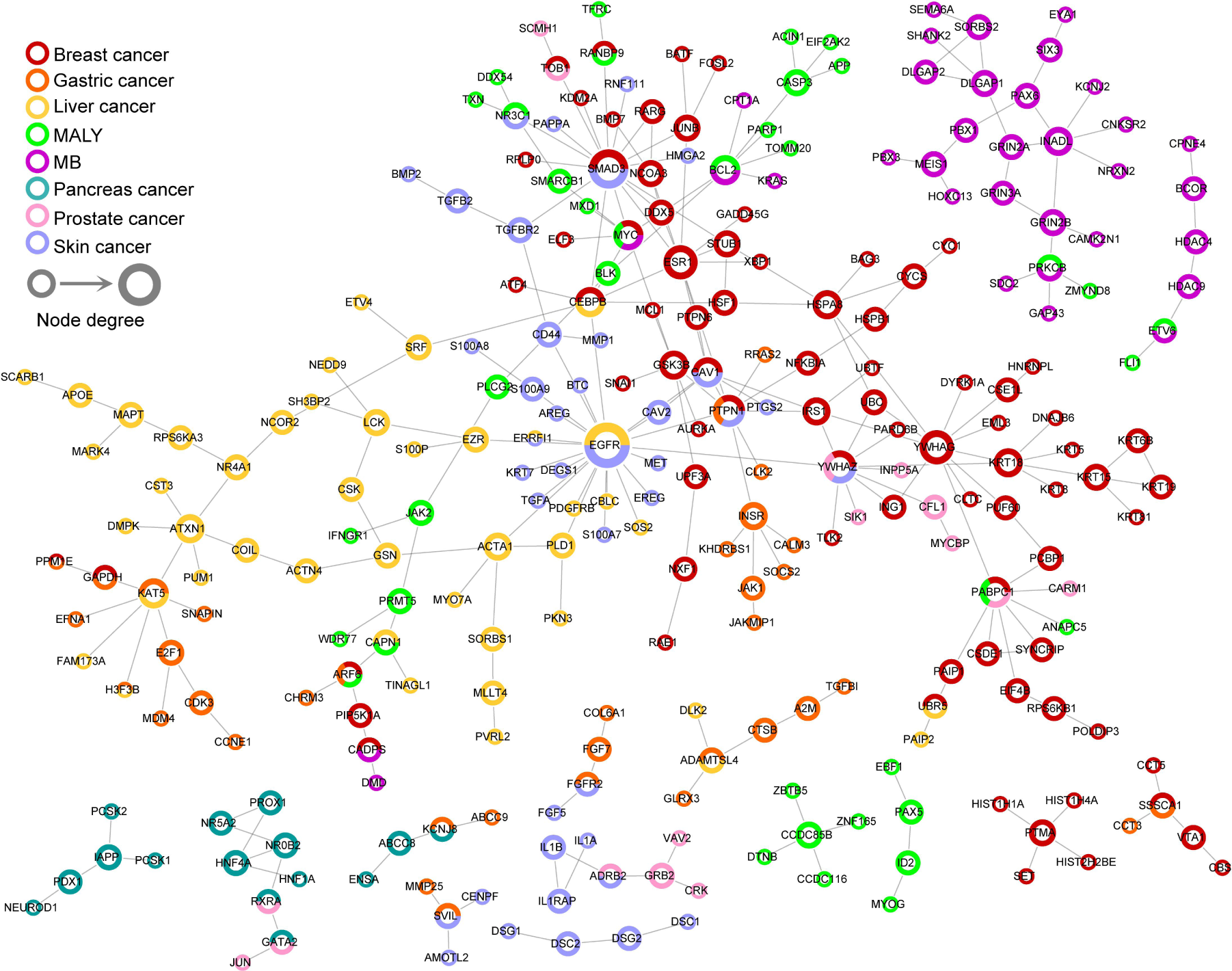
Overview of SME-networks identified by Ennet. Subnetworks with node number > 3 are shown.

Most SME-networks found by Ennet were very specific, the vast majority (92.6%, 404/436) of the genes being unique to only one cancer type and no gene occurring in SME-networks of 4 or more cancer types (Fig. 3A). This suggests that tumors from different tissues have different driver events. To further investigate which pathways were enriched for Ennet detected genes, we performed a KEGG pathway enrichment analysis. Pathways with significant enrichment of Ennet genes were also largely cancer type-specific (Fig. 3B). For example, the two pathways most enriched with genes detected in breast cancer were the genes belong to the “hippo signaling pathway” and the “breast cancer” pathway, and in liver cancer they were genes belonging to the “MAPK signaling pathway” and “regulation of actin cytoskeleton” pathways. In addition, several pathways were shared among different cancer types, like the PI3K-Akt signaling pathway, the MAPK signaling pathway and the ErbB signaling pathway. Taken together, genes identified by Ennet are highly cancer type-specific.

**Figure 3.**
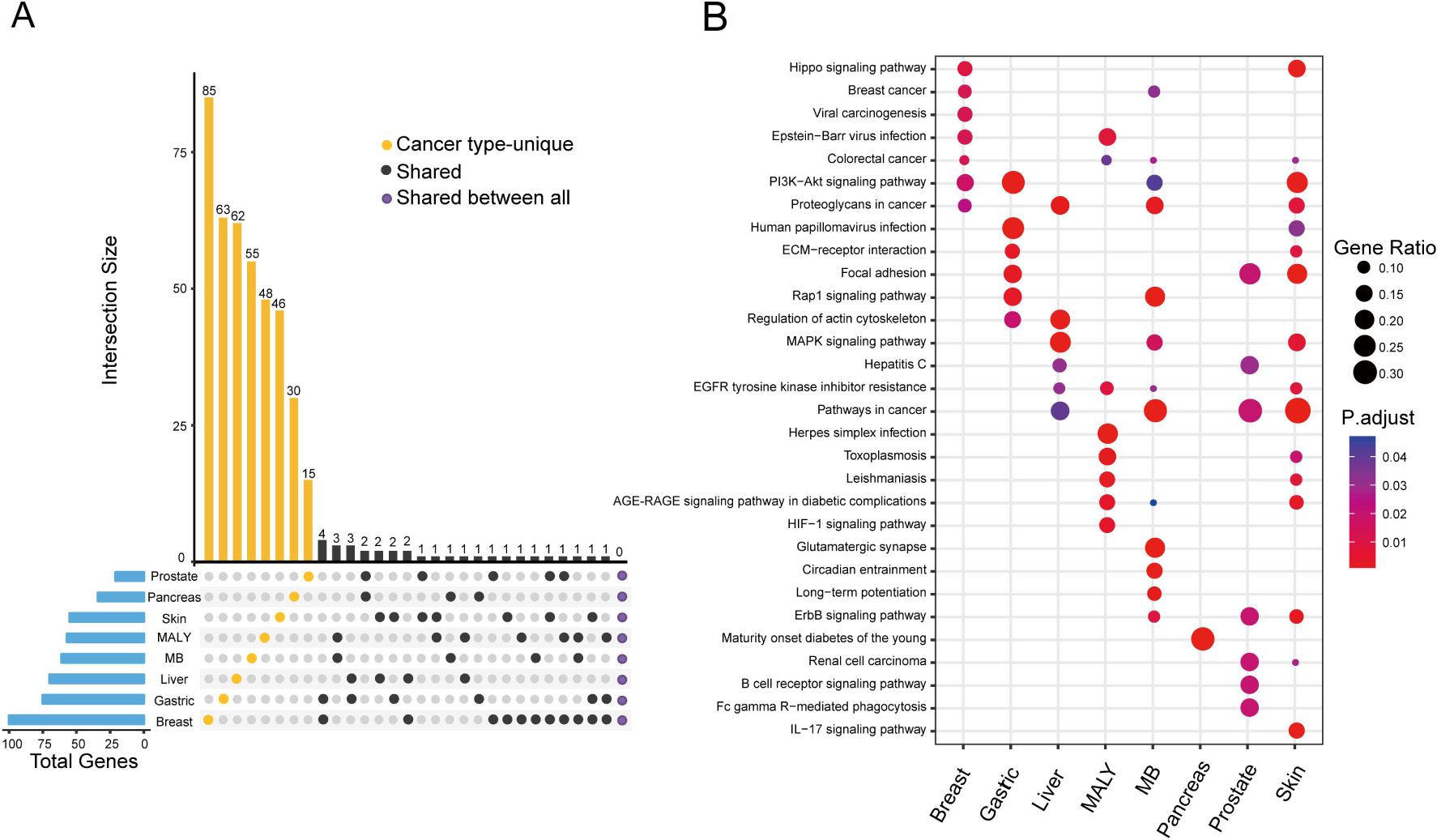
Genes detected by Ennet were cancer related and cancer type-specific. (A) KEGG enrichment analysis for genes in SME-networks identified by Ennet. (B) Numbers of genes identified by Ennet in 8 cancers.

### Genes in SME-networks have few mutations in the gene bodies

Currently, approaches for discovering driver genes mainly relied on mutational information from the gene body regions. In contrast, Ennet uses mutations within enhancer elements, which are typically ignored or unutilized, rather than mutations in gene bodies. To illustrate the differences between Ennet and conventional driver hunters, we compared the genes in the SME-networks and genes predicted by MutSigCV [25], a state-of-the-art cancer driver caller. MutSigCV and Ennet show few overlaps in their predictions within all 8 cancer types (Fig 4A). We next investigated the mutational spectrum of the genes identified by the two methods and found large divergences (malignant lymphoma (MALY) is shown in Fig. 4B, other cancers can be found in Supplemental Figure 2 to Supplemental Figure 8). In MALY, genes detected by Ennet all have high mutation rate in their enhancers (from 5.0% to 54.6%), and low mutation rates in their exons (only two genes had mutation rates above 20%, *MYC* and *BCL2*). Yet, those found by MutSigCV all have high exon mutation rate (from 3.5% to 40.4%), and have low mutation rates in their distal regions (0%-14.9%, and in 9/34 genes mutations were only found in the exons). This indicates that genes in SME-networks have few mutations in gene bodies. Next, we compared the genes discovered in MALY versus genes in the Cancer Gene Census (CGC) [26] and found 9 genes with a strong support of being cancer driver genes in CGC (Fig. 4B), suggesting that the other 48 genes may also function in cancer. For example, *CASP3*, an executioner caspase, plays a key role in the execution-phase of cell apoptosis [27] and high levels of uncleaved caspase 3 were associated with decreased in acute myelogenous leukemia [28]. Therefore, genes detected by Ennet and MutSigCV are intrinsically different in the mutational spectrum and Ennet can serve to complement existing methods.

**Figure 4.**
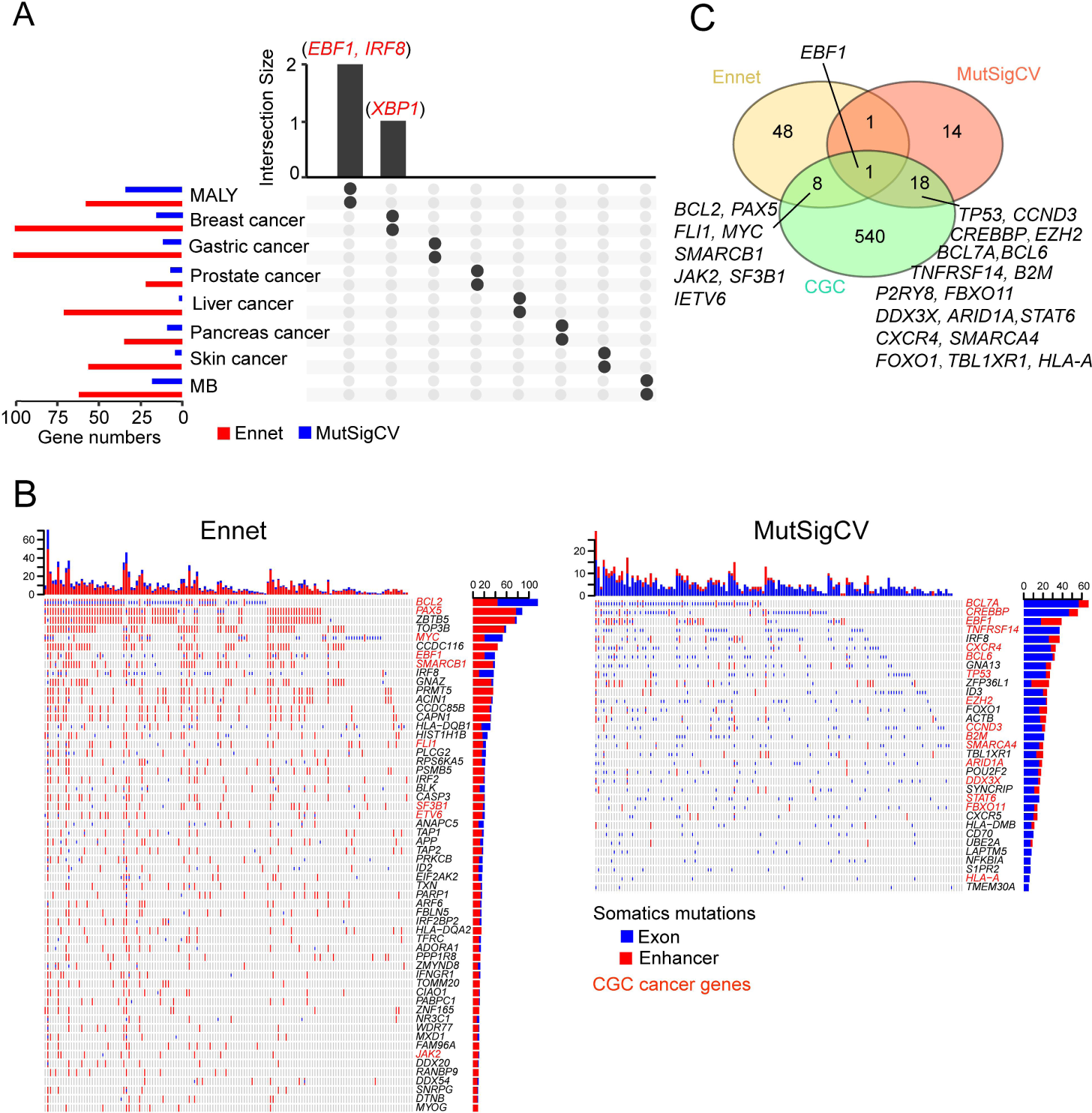
Genes in SME-networks have few mutations in the gene bodies. (A) The overlap of genes identified by Ennet and MutSigCV across different cancer types. Few genes were detected simultaneously by both methods. (B) Mutation profiles of genes found by Ennet (left) and by MutSigCV (right) in malignant lymphoma (MALY). Almost all genes identified by Ennet have few lesions in their exon regions. (C) Venn diagram of genes in (B) and cancer genes in CGC.

### Ennet is more efficient than mutation density-based methods

The network-based method employed by Ennet combines mutational information from multiple enhancers and may be more efficient in exploration of SMEs than density-based methods (Fig. 5A). A recent study analyzed the mutations of 150 chronic lymphocytic leukaemia (CLL) WGS samples in an unbiased fashion using a mutation density-based method, and showed that most of the hotspots were in the coding regions and in known target elements of the somatic hypermutation (SHM) process [5]. Through similar methods, we identified only one hotspot in the enhancer of *PAX5*, also observed by Puente *et al.* (Fig. 5B). We then applied Ennet to the same dataset. Although the enhancers of most genes (except those affected by the SHM process) had low mutation frequencies (Fig. 5C), Ennet successfully identified 38 genes in 8 SMEs enriched subnetworks, of which 3 subnetworks contained more than 3 genes (Fig. 5D). The subnetwork containing *PAX5* included three additional genes, *EBF1*, *ID2* and *TCF3*. Early B-cell factor 1 (*Ebf1*) is a transcription factor with documented dose-dependent functions in normal and malignant B-lymphocyte development [29], and a synergistic effect of combined dose reduction of *EBF1* and *PAX5* was observed in mice [30]. The interplay between *PAX5* and *ID2*, *TCF3* in B cell lineage commitment and leukemia was widely reported [31–33]. These suggest that these three genes have been ignored by density-based methods, may also promote CLL development. Besides, many well-known cancer genes in CLL such as *BCL2*, *MYC*, *BLK* and *LYN* were included in the SME-networks detected by Ennet (Fig. 5D). Collectively, Ennet is more efficient in exploration of SMEs than mutation density-based methods.

**Figure 5.**
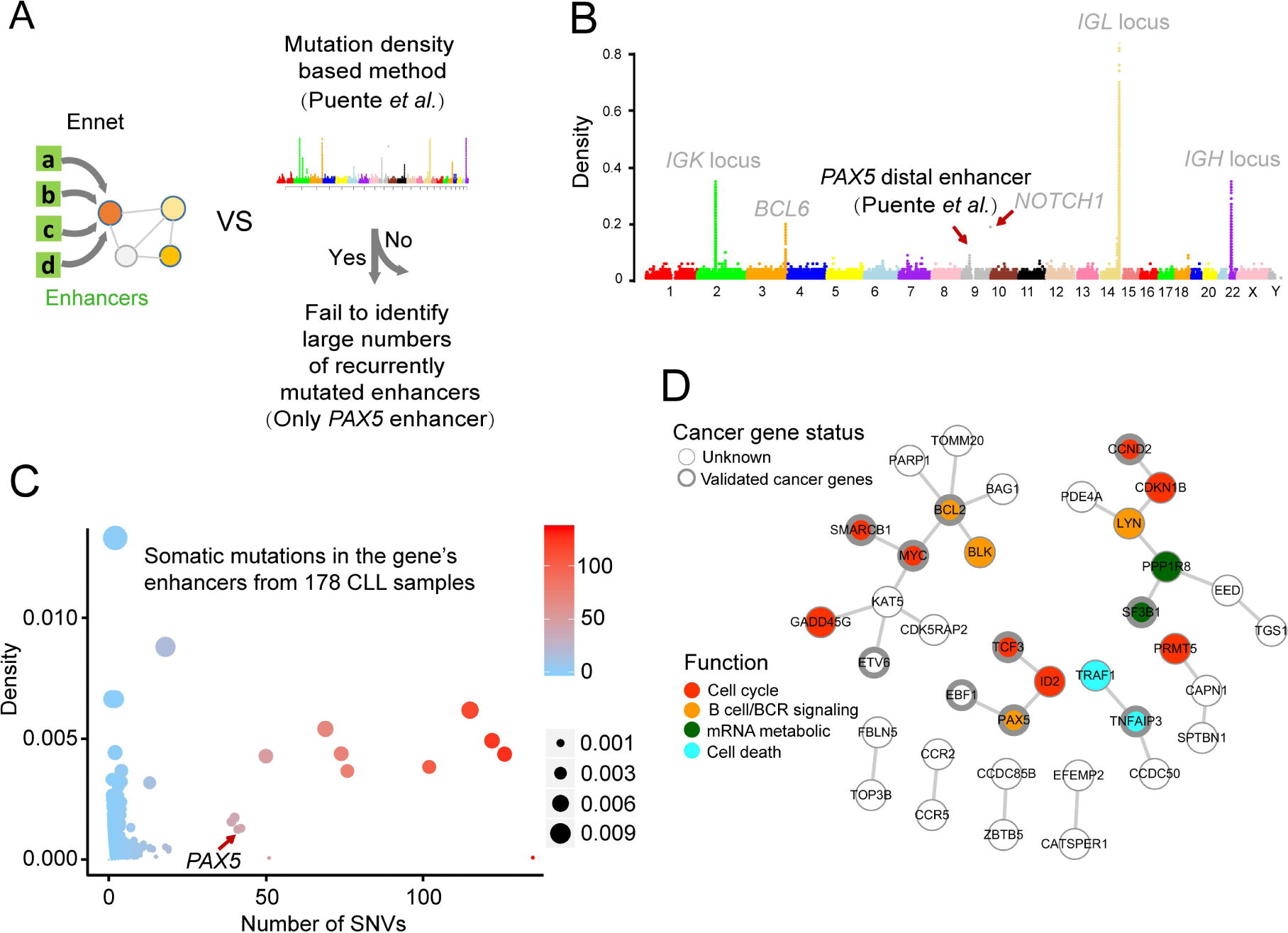
Ennet is more efficient than mutation density-based methods. (A) Schematic of the comparison between Ennet and mutation density-based methods. (B) Regions with a high density of somatic mutations in 178 CLL samples. Regions correspond to recurrently mutated coding genes (grey), targets of SHM (grey), and enhancers (black). (C) Somatic mutation density of genes’ enhancers from 178 CLL samples. (D) SME-networks identified by Ennet in CLL.

### Ennet identified a novel potential cancer-driving network in MB

MB is the most common malignant brain tumor of children [34]. We noticed that most of the genes in the biggest SME-network of MB have less well characterized roles in MB (Fig. 6A). When we searched PubMed using gene/protein name and medulloblastoma as keywords, relevant records were only found for *PAX6* (4 records), and *EYA1* (1 record) (by November 2017). Pathway analysis for genes in the biggest SME-network of MB demonstrated that these genes were enriched in glutamate receptor signaling and neural development associated pathways (Fig. 6B). Dysregulation of developmental pathways can be an important mechanism underlying cancer development [35]. Genes in the glutamate receptor signaling pathway, including *GRIN2A*, *GRIN3A*, and *GRIN2B*, are significantly down-regulated in the glioblastoma than in matched normal tissue (Supplemental Figure 9). Taken together, our results suggest that genes dysregulated in these SME-networks may be involved in the development of MB.

**Figure 6.**
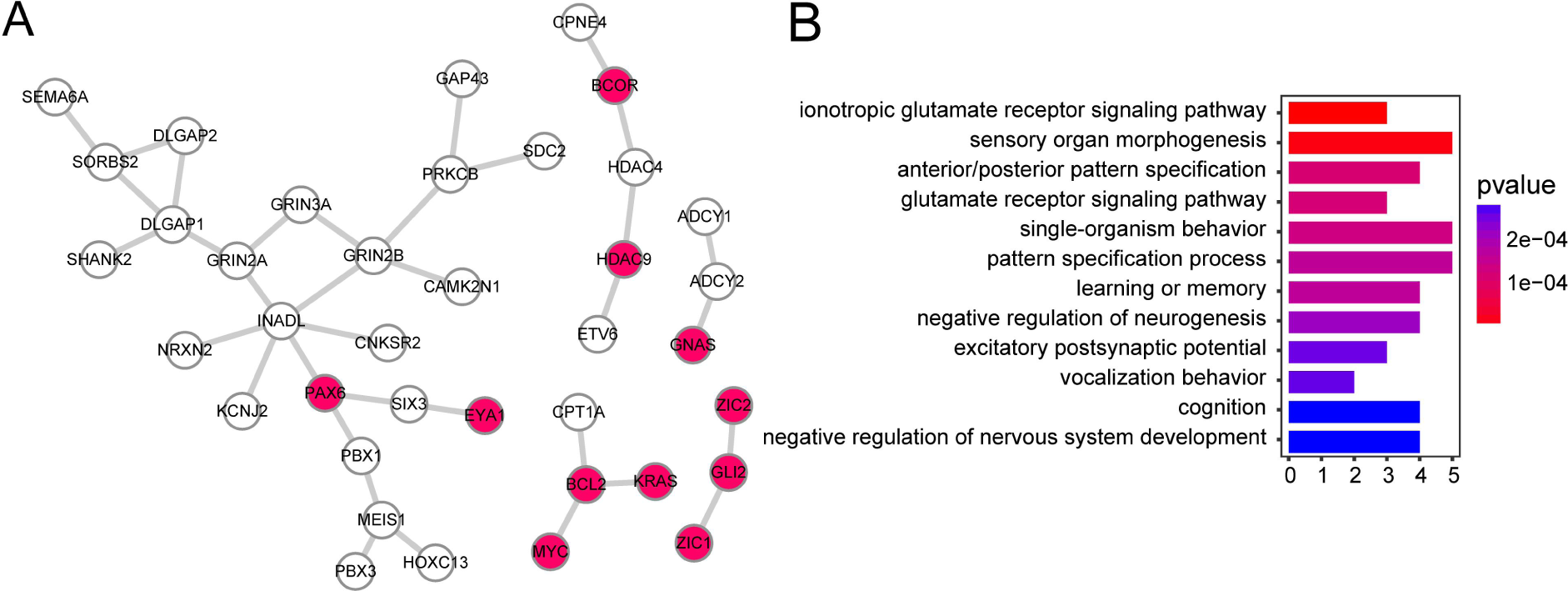
Ennet found a potentially new pathway involved in MB. (A) SME-networks identified by Ennet in MB. Red nodes indicate that there were relevant records in the PubMed database. (B) Genes in the biggest SME-networks were enriched in glutamate receptor signaling and neural development pathways.

## DISCUSSION

Whole-genome sequencing provides us with a large number of, but very sparse, somatic mutations in enhancers. Over the past few years, we have learned very little about cancer from such mutations. Several interesting questions can be asked concerning these mutations. Can we get more information about cancers by a focused analysis of these sparse enhancer mutations? Are there any genes or pathways in which important mutations occur in the enhancers rather than in the gene body or promoter regions?

In this study, we designed Ennet to answer these questions. We applied Ennet to analyze somatic mutations in enhancers across 8 cancer types through integrating enhancer-gene interactions with knowledge-based gene interaction networks. We were surprised to find that these unutilized enhancer-only somatic mutations enabled us to identify many well-known cancer driver genes, like *ESR1* in breast cancer [3] and *PAX5* in CLL [5]. Notably, almost none of the genes identified by Ennet, whether known or unknown, were significantly mutated in gene body or promoter region (identified by MutSigCV, q value < 0.05). Moreover, Ennet identified a potentially novel cancer-driving biological network in MB. In summary, our work suggested that using only somatic mutations in enhancers can be an effective way to discover potential cancer-driving biological networks and thus Ennet provides a new way to investigate SMEs in tumor development.

Information on enhancer-gene interactions are critical for linking enhancers to knowledge-based networks in Ennet. Although we have integrated enhancer-gene pairs from various sources including both high-throughput experiments and computational prediction, the interaction maps may still be incomplete. Hi-C, ChIA-PET and similar experiments [36] have provided a mass of enhancer-gene pairs, but thus far these data are only available for a few human cell types. Several cancer types such as ovary cancer are not suited to analysis by Ennet owing to lack of interaction maps in the associated tissues. Fortunately, work such as the 4D Nucleome program [37] will continue to yield information about the chromosome architecture in diverse cell types in the near future. With the availability of increasing number of whole cancer genomes and a better understanding of 3D chromatin structure in different cell types, the exploration the role of mutations in regulatory regions has just begun. We expect that Ennet will serve as a complementary method to other cancer driver callers and may lead to new insights in cancer genomic research.

## METHODS

### Acquisition whole genome somatic mutation data

We obtained precomputed somatic variants from the whole-genome sequences of 1,648 samples across 7 cancer types from the ICGC database (Release 25) [38], 256 samples of 4 cancer types from Alexandrov *et al*. [39], and 100 gastric cancer samples from the supplemental materials of Wang *et al*. [40]. All data from ICGC should be free of a publication moratorium according to the ICGC Publication Policy (<u>http://docs.icgc.org/portal/publication/</u>). All samples in the PanCancer Analysis of Whole Genomes (PCAWG) project were excluded because of usage limitation. For duplicate samples of ICGC database and Alexandrov *et al*., only data from ICGC was kept. Finally, 8 cancer types were included for the subsequent analysis: breast cancer, gastric cancer, liver cancer, malignant lymphoma (MALY), medulloblastoma (MB), pancreas cancer, prostate cancer, and skin cancer (Fig. 1B and Supplemental Table 1).

### Collecting and processing enhancer-promoter interactions

We collected enhancer-promoter interactions from two sources: 1) Chromatin interactome from high throughput experiments, such as Capture Hi-C. 2) Predictions made by six computational methods: RIPPLE [17], IM-PET [16], PreSTIGE [15], Epitensor [18], PETModule [19], and JEME [20]. Supplemental Table 2 listed all the sources and annotations used in this study. Except for GM12878 cell line with Capture Hi-C data [41], interactions of specific cell type from different studies were merged. Then we performed the filtering on the putative distal regulatory elements (DREs). For a given DRE, a 1bp overlap with enhancer-like regions annotated by ENCODE was required [1] if the corresponding enhancer annotations was available. In order not to be affected by coding regions, any DRE overlapping with exons of protein-coding genes were excluded. The annotation we used were GENCODE v24 downloaded from GENCODE’s website [42]. Enhancers function by binding with transcription factors. We therefore divided the DRE region according to transcription factor binding data generated by ENCODE if the corresponding TF binding data was available. The manipulation on genomic features were done using BEDTools [43] and BEDOPS [44]. The integrated enhancer-gene interactions are available at: http://bigdata.ibp.ac.cn/ennet/.

### Knowledge-based gene networks

In principle, knowledge-based gene networks can be based on any type of interactions between genes. In this version, Ennet collects protein-protein interaction networks (HPRD [45], iRefIndex [46] and Multinet [47]) and KEGG pathways [48].

### Integrated analysis with Ennet

Three key inputs are required for Ennet: whole genome somatic mutations, enhancer-gene interactions and knowledge-based gene networks. Ennet begins by counting noncoding mutations falling within tissue-specific enhancers for each gene. It then sums up all mutations across patients and maps the count to a knowledge based gene network. After mapping, a random walk based network propagation [8,49] is applied to ‘smooth’ the mutation signal across the network (this step is optional). Finally, Ennet selects utilize a python package NetworkX [50] to select subnetworks.

### Cancer driver gene and cancer signaling pathway data

We collected the cancer driver genes from the widely used Cancer Gene Census (CGC, http://cancer.sanger.ac.uk/census) [26] and from the Network of Cancer Genes database (version 4) [51]. In addition, we also collected genes from cancer signaling pathways annotated in the Netpath database [52].

### Other bioinformatic analysis

UpSet plots in Figure 3 and Figure 4 were generated by R package UpSetR [53]. Waterfall plots in Figure 4 and Supplemental Figure 2-8 were generated using R package ComplexHeatmap v1.14.0 [54]. Enrichment analysis was done by R package clusterProfiler [55]. For the comparison with MutSigCV [25], we first annotated the mutations using Oncotator v1.9.3.0 [56] and then feed MutSigCV v1.4.1 with MAF files following the instructions (http://archive.broadinstitute.org/cancer/cga/mutsig).

## Acknowledgements

We thank Dr. Geir Skogerbø for careful reading and valuable suggestions on the manuscript and the staff of the Protein Research Core Facility at the Institute of Biophysics, Chinese Academy of Sciences, Dr. Xiaowei Chen and Dr. Zhen Fan, for their assistance in highthroughput sequencing data analysis and discussion. Data analysis and computing resource was supported by Center of Big Data Research in Health (http://bigdata.ibp.ac.cn), Institute of Biophysics, Chinese Academy of Sciences.

## Author Contributions

Ya Cui, Runsheng Chen, Yiwei Niu and Xueyi Teng conceived the project; Ya Cui, Yiwei Niu and Xueyi Teng developed the method; All authors performed the data analysis; All authors drafted the manuscript and approved the final version.

